# Gene losses in the common vampire bat illuminate molecular adaptations to blood feeding

**DOI:** 10.1101/2021.10.18.462363

**Authors:** Moritz Blumer, Tom Brown, Mariella Bontempo Freitas, Ana Luiza Destro, Juraci A. Oliveira, Ariadna Morales, Tilman Schell, Carola Greve, Martin Pippel, David Jebb, Nikolai Hecker, Alexis-Walid Ahmed, Bogdan Kirilenko, Maddy Foote, Axel Janke, Burton K. Lim, Michael Hiller

## Abstract

Feeding exclusively on blood, vampire bats represent the only obligate sanguivorous lineage among mammals. To uncover genomic changes associated with adaptations to this unique dietary specialization, we generated a new haplotype-resolved reference-quality genome of the common vampire bat (*Desmodus rotundus*) and screened 26 bat species for genes that were specifically lost in the vampire bat lineage. We discovered previously-unknown gene losses that relate to metabolic and physiological changes, such as reduced insulin secretion (*FFAR1*, *SLC30A8*), limited glycogen stores (*PPP1R3E*), and a distinct gastric physiology (*CTSE*). Other gene losses likely reflect the biased nutrient composition (*ERN2*, *CTRL*) and distinct pathogen diversity of blood (*RNASE7*). Interestingly, the loss of *REP15* likely helped vampire bats to adapt to high dietary iron levels by enhancing iron excretion and the loss of the 24S-hydroxycholesterol metabolizing enzyme *CYP39A1* could contribute to their exceptional cognitive abilities. Finally, losses of key cone phototransduction genes (*PDE6H*, *PDE6C*) suggest that these strictly-nocturnal bats completely lack cone-based vision. These findings enhance our understanding of vampire bat biology and the genomic underpinnings of adaptations to sanguivory.

## Introduction

Vampire bats are the only obligate sanguivorous lineage among tetrapods ^1^. This exceptional dietary specialization is reflected in all aspects of their biology, including morphology, physiology and behavior ^2,3^. To detect prey, the common vampire bat (*Desmodus rotundus*) exhibits a well-developed olfactory system ^4^, advanced low-frequency hearing abilities ^5^ and, unique among mammals, the ability to sense infrared radiation ^6^. Compared to other bats, vampire bats have exceptional terrestrial locomotion skills to sneak up on their prey ^7^. Razor-sharp enamel-less upper incisors help cutting through the prey’s skin and anticoagulants in their saliva prevent the prey’s blood from coagulating during feeding ^8,9^.

As the sole nutritional source, blood represents a challenging diet for several reasons. First, blood has a high fluid content of 78% and a comparatively low caloric value, making it necessary for a vampire bat to ingest as much as 1.4 times their body weight in blood during a single meal ^10,11^. To enable the ingestion of large amounts of blood, their stomach experienced a functional shift towards a distensible organ primarily engaged in storage and fluid absorption ^12^. Second, blood has a high iron content compared to other diets, which mainly stems from hemoglobin-derived heme and ferric iron transported by transferrin ^13–15^. Third, the dry mass of blood has a highly skewed nutritional composition, providing mostly proteins (93%) with very little lipids and carbohydrates (1% each) ^11^.

Due to a low carbohydrate intake, vampire bats exhibit lower basal insulin levels than other mammals ^10,11,16^. Similar to human type 2 diabetes patients, vampire bats feature a reduced glucose-stimulated insulin secretion response, resulting in hyperglycemia upon an experimental glucose overload ^16^. Glycogen and lipid stores are also reduced in vampire bats, which contributes to their fasting vulnerability and early deaths after 48-72 h of fasting ^17,18^. To compensate for their fasting vulnerability, vampire bats share regurgitated blood with roost mates that failed to obtain a nightly meal ^19,20^.

To gain insights into the molecular basis of vampire bat adaptations to sanguivory, comparative studies using the genome of the common vampire bat detected signatures of selection in genes involved in responses to nutrient starvation, metabolism, nitrogen waste disposal, the coagulation cascade, and immunity ^21,22^. Common vampire bats have fewer *TAS2R* taste receptor genes than other bats, indicating a reduced sense of bitter taste reception ^23,24^. In addition to genetic changes, vampire bats also possess a gut microbiome very different from that of other bats, and genes encoded by their gut microbiome further contribute to meeting the challenges of sanguivory ^21^. Despite these advances, our understanding of which genomic changes are important for adaptations to sanguivory remains incomplete.

A limitation of previous comparative studies was the restricted taxonomic representation. For example, Zepeda Mendoza *et al*. ^21^ compared the *D. rotundus* genome to nine other bat genomes, with *D. rotundus* as the only representative of Phyllostomidae bats (Figure 1A). With this limited taxonomic resolution one cannot differentiate between those genomic changes that are shared between all phyllostomid bats (comprising more than 200 species) and those changes that evolved specifically in the vampire bat lineage and thus could be relevant for adaptations to sanguivory. To uncover genomic changes that evolved specifically in the vampire bat lineage, we generated a state-of-the-art haplotype-resolved chromosome-level assembly of *D. rotundus* and made use of many additional phyllostomid genomes that include the closest sister species of vampire bats (Supplementary Table 1). Since the loss of ancestral genes can be an important evolutionary force and previous studies revealed many associations between gene losses and phenotypic differences, including dietary adaptations ^25–29^, we conducted a genome-wide screen for genes that are specifically lost in *D. rotundus* and that lack inactivating mutations in 25 other bat species. This screen revealed 3 known and 10 previously-unknown gene losses. Many of these novel gene losses have clear associations with vampire bat traits and some gene losses may contribute to coping with the challenges imposed by sanguivory.

**Figure 1:**
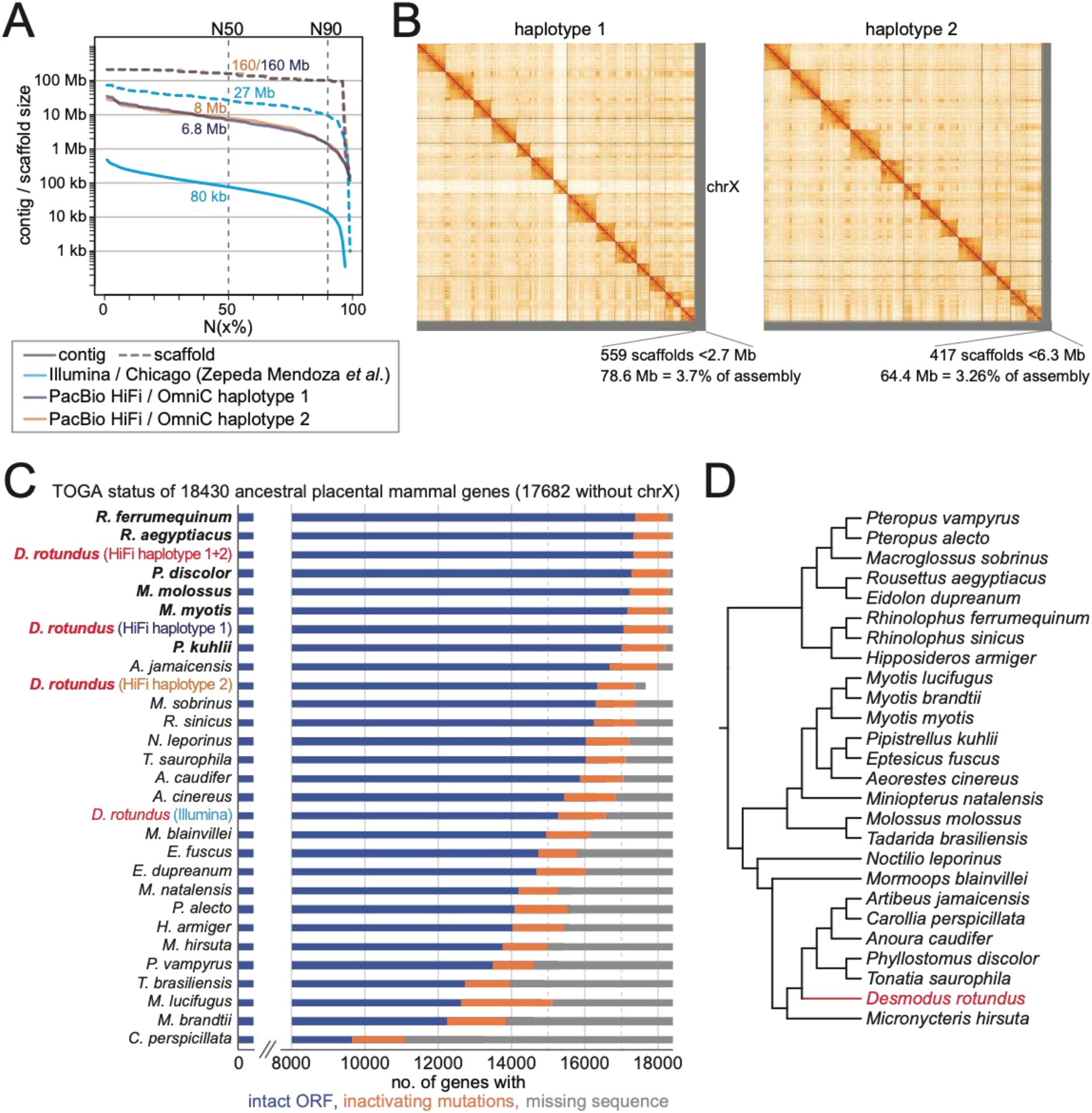
Haplotype-resolved chromosome-level assembly of *Desmodus rotundus*. (A) Comparison of assembly contiguity between our haplotype-resolved assemblies and the previous Illumina-based assembly ^21^. The graph shows contig (solid lines) and scaffold (dashed lines) sizes at the Y-axis, for which x per cent of the assembly consists of contigs and scaffolds of at least that size. (B) HiC contact maps show 14 and 13 chromosome-level scaffolds for haplotype 1 and 2 that each comprise more than 96% of the respective haplotype assembly. Haplotype 1 contains the X chromosome. The Y chromosome is part of haplotype 2 and assembled into several shorter scaffolds. (C) Status of 18,430 ancestral placental mammal genes in each assembly, as inferred by TOGA. Genes are classified into those that have an intact reading frame (blue), have inactivating mutations (orange) or have missing coding parts due to assembly gaps or fragmentation (grey). Assemblies are sorted by the number of intact genes. Long readbased assemblies (bold font) consistently exhibit more intact genes and fewer missing genes compared to short readbased assemblies (not bold). For the *D. rotundus* haplotype 2 assembly that does not contain the X chromosome, we excluded genes located on the X chromosome and only considered the remaining 17,682 genes. To provide a fair comparison with previous assemblies that collapse both haplotypes, we also computed statistics for the union of both *D. rotundus* haplotype assemblies (haplotype 1+2), which exhibits the third highest number of intact genes of all included genomes. (D) Phylogeny of the bats analyzed in this study ^1,42,43^.

## Results

### A new haplotype-resolved reference-quality *D. rotundus* assembly

To perform an accurate and comprehensive screen for gene losses in the common vampire bat, a genome assembly with high completeness, contiguity and base accuracy is desirable. The existing Illumina assembly of the common vampire bat ^21^ has more than 50,000 assembly gaps, indicating that a subset of genes will have missing sequences. This assembly also exhibits base errors that mimic gene-inactivating mutations, including cases where multiple putative mutations in the same gene are all erroneous (Supplementary Figure 1). Therefore, we generated a new reference-quality *D. rotundus* assembly. We used PacBio circular consensus (HiFi) sequencing ^30^ to produce 32X coverage in reads with an average length of 9.1 kb and used the Dovetail Omni-C protocol to produce 67X coverage of chromosome conformation capture Illumina read pairs. With these data, we obtained two haplotype-resolved chromosome-level assemblies that have only 596 and 557 assembly gaps, contig N50 values of 6.85 and 7.98 Mb and scaffold N50 values of 160.1 and 160.1 Mb (Figure 1A, Supplementary Table 1). The contig N50 metric is ~90-times higher than that of the Illumina-based *D. rotundus* assembly ^21^. The karyotype of *D. rotundus* is 2n=28 ^31^. All 13 autosome pairs are represented by chromosome-level scaffolds (Figure 1B). Haplotype 1 contains the X chromosome as an additional chromosome-level scaffold. Haplotype 2 contains the Y chromosome, which was assembled as several scaffolds. Overall, more than 96% of both haplotypes are contained in chromosome-level scaffolds (Figure 1B). Using Merqury ^32^, we estimated a very high base accuracy of QV 64.2 and 64.5 for both haplotypes, indicating one error per ~2.6 million base pairs.

To systematically assess gene completeness among the *D. rotundus* and other available bat genomes, we applied TOGA (Tool to infer Orthologs from Genome Alignments), a method that infers orthologs from whole genome alignments ^33^, to 18,430 ancestral placental mammal coding genes (Methods). In comparison to the Illumina *D. rotundus* assembly, the number of ancestral genes with missing sequences dropped from 1,841 to 128 in our combined haplotype assemblies, whereas the number of genes with an intact reading frame increased from 15,295 to 17,301 (Supplementary Table 1). This indicates a substantially higher gene completeness in our assemblies, similar to the most contiguous bat assemblies generated to date ^34^ (Figure 1C).

### A genome-wide screen revealed known and novel vampire bat-specific gene losses

To detect genes that were specifically lost in the vampire bat lineage, we used our two haplotype assemblies and considered 25 other bat genomes ^21,23,34–41^ (http://dnazoo.org/), including *Micronycteris hirsuta* and *Phyllostomus discolor*, which represent the closest phylogenetic outgroup and sister lineages of vampire bats, respectively ^1,42,43^ (Figure 1D). Using the human gene annotation as the reference, we applied TOGA to detect genes that exhibit gene-inactivating mutations (premature stop codons, frameshifts, splice site disruptions, and deletions of exons or entire genes) specifically in *D. rotundus*.

This screen revealed 13 vampire bat-specific gene losses. Three of these losses have been reported before: the sweet taste receptor gene *TAS1R2* ^44^ and the bitter taste receptor genes *TAS2R5* and *TAS2R42* ^24^. These gene losses indicate a reduced sense of taste reception in vampire bats. To our knowledge, the remaining 10 gene losses (*REP15*, *FFAR1*, *SLC30A8*, *PPP1R3E*, *CTSE*, *ERN2*, *CTRL*, *CYP39A1*, *PDE6H*, *RNASE7*) have not been reported before.

The inactivating mutations in these 10 genes are displayed in Figure 2A. In support of the validity of the underlying mutations, the 10 gene losses are also detected in the Illumina *D. rotundus* assembly and exhibit at least one shared inactivating mutation that is supported by both PacBio HiFi and Illumina reads. Further supporting gene loss, selection rate analysis using RELAX ^45^ showed that 9 of the 10 genes evolve under relaxed selection in *D. rotundus* to preserve the reading frame (significant for 7 of 9 genes, Supplementary Table 2). Finally, inspecting available *D. rotundus* RNA-seq data, we either found no relevant expression in tissues where expression would be expected or found that the aligned reads supported the gene-inactivating mutations, indicating that potential transcripts cannot be translated into a full length protein (Supplementary Figure 2).

**Figure 2:**
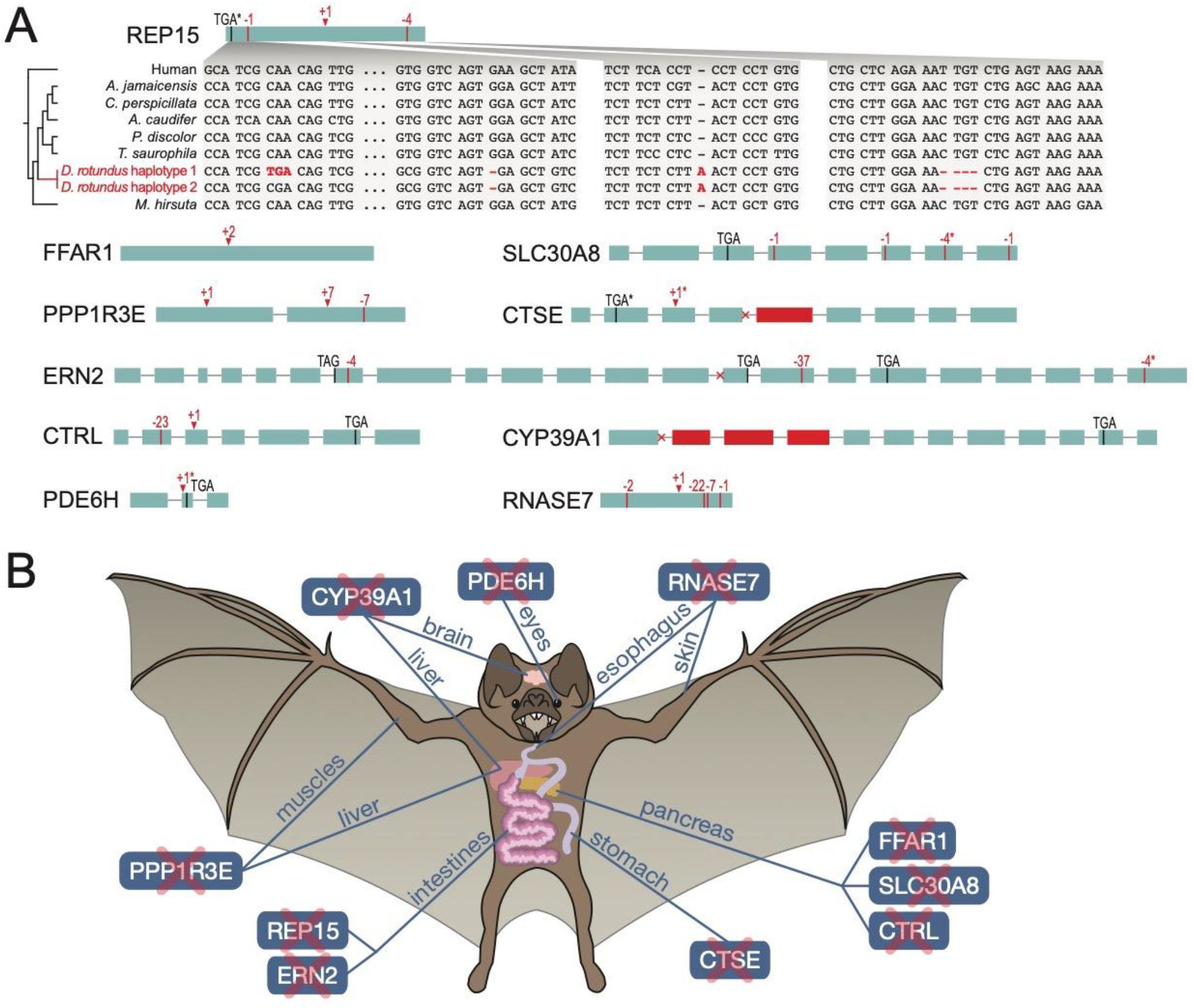
Inactivating mutations and affected organ systems of 10 previously-unknown vampire bat-specific gene losses. (A) Exon-intron structure visualization with inactivating mutations that are detected in the *Desmodus rotundus* genome. Premature stop codons are shown as black vertical lines, frameshifting deletions as red vertical lines and frameshifting insertions as red arrow heads. Donor or acceptor splice site mutations are indicated as a cross at the exon boundaries. Deleted exons are in red. Asterisks denote mutations that are heterozygous in our sequenced *D. rotundus* individual (present in only one of the two haplotype assemblies). The inset for *REP15* illustrates that inactivating mutations and thus gene loss was only detected in the common vampire bat. (B) Illustration of organs and anatomical sites where the 10 genes play important roles.

Intriguingly, as detailed below, the loss of these genes in the common vampire bat relates to a number of adaptations to their highly specialized blood diet, such as reduced insulin secretion and glycogen synthesis and a distinct stomach physiology (Figure 2B). These gene losses further provide a possible mechanism contributing to vampire bats’ exceptional social behavior, and indicate the complete lack of cone photoreceptor function in *D. rotundus*.

### Loss of *REP15* and enhanced iron excretion

The loss of *REP15* (RAB15 effector protein), a gene involved in regulating cellular iron uptake ^46^, is likely related to the obligatory iron-rich blood diet of vampire bats. Despite the importance of iron for various cellular processes, iron overload can have severe detrimental effects ^15^. Remarkably, the common vampire bat tolerates extreme dietary iron levels without exhibiting adverse effects ^47^ – the relative amount of dietary iron was estimated to be 800-fold higher compared to humans ^13^. However, blood iron concentration in vampire bats has never been determined to our knowledge. Therefore, we measured whole blood iron concentrations in wild *D. rotundus* and compared it to two other neotropical bats (Supplementary Table 3). We found that blood iron content was significantly higher in the common vampire bat compared to the frui-teating bat *Artibeus lituratus* (family Phyllostomidae; t-test: P=0.01, Figure 3A). Vampire bats also had higher blood iron levels compared to the insectivorous outgroup bat *Myotis myotis* (family Vespertilionidae; not significant, likely due to the small sample size of four individuals; Figure 3A). This suggests that vampire bats would benefit from mechanisms to lower systemic iron levels.

**Figure 3:**
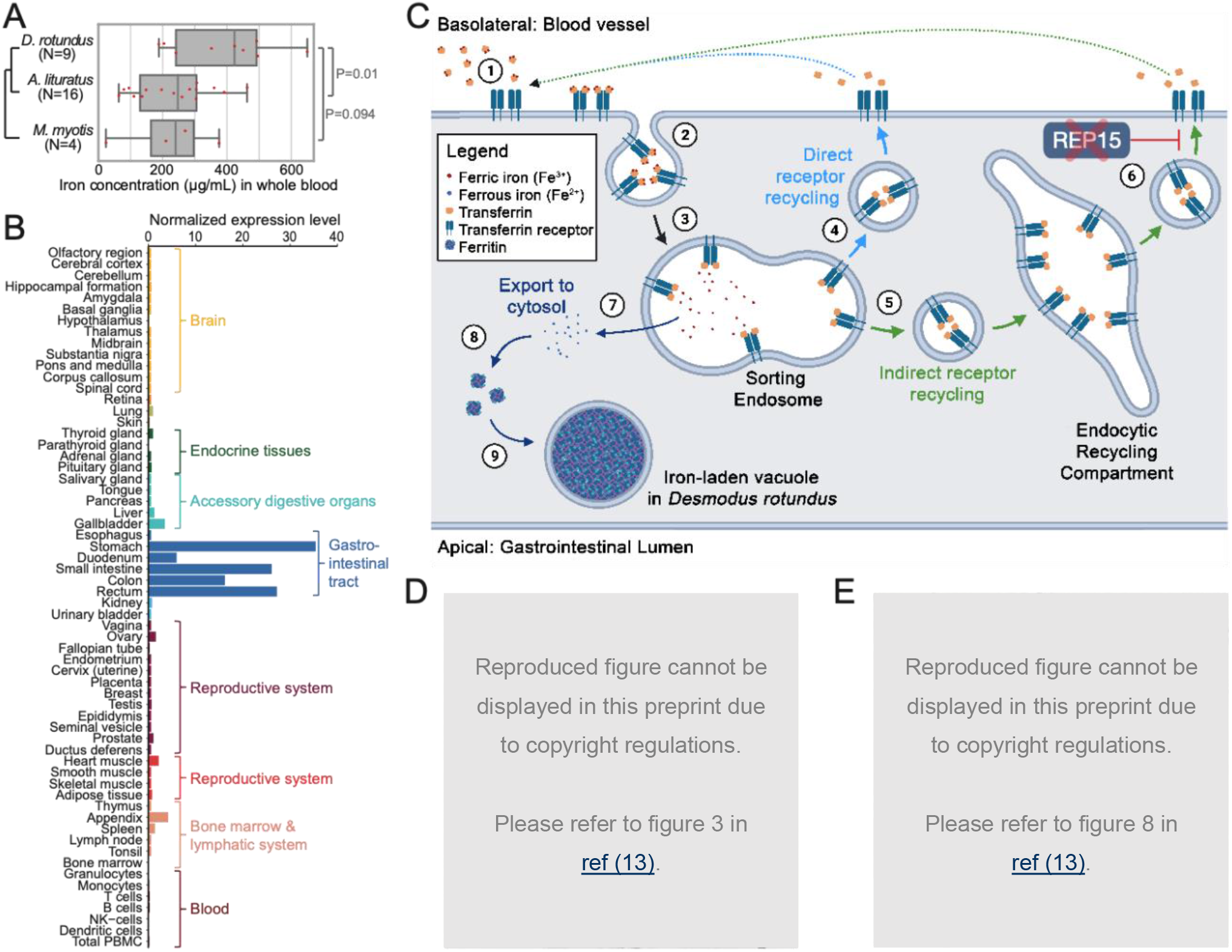
Loss of *REP15* in *Desmodus rotundus* and enhanced iron excretion. (A) Measurements of iron concentration in whole blood of *D. rotundus, Artibeus lituratus* and *Myotis myotis* show that vampire bats have higher circulatory iron levels. P-values are computed with a two-sided t-test. (B) *REP15* mRNA expression is highest in gastrointestinal tract tissues. Data was taken from the Human Protein Atlas ^48^ (https://www.proteinatlas.org/ENSG00000174236-REP15/tissue) and show the consensus RNA expression values that integrate three gene expression datasets. (C) Illustration of transferrin receptor-mediated cellular iron uptake and function of *REP15* in intestinal epithelial cells. Transferrin, an abundant ferric iron-binding plasma protein, binds to transferrin receptors that are present only in the basolateral membrane (1). Transferrin-transferrin receptor complexes are internalized via endocytosis (2). In sorting endosomes, ferric iron is released (3) and the unladen complexes are either directly targeted back to the cell membrane (4) or sent to the endocytic recycling compartment (5). REP15, encoded by the gene that is lost in *D. rotundus*, specifically localizes to the endocytic recycling compartment (6) and inhibits recycling of the unladen complex to the cell membrane ^46^, where transferrin and its receptor dissociate and the released transferrin can bind ferric iron again. Because the availability of transferrin receptors on the cell surface limits iron uptake ^49^, the presence of *REP15* normally inhibits cellular iron uptake. In the sorting endosome, ferric iron (Fe^3+^) is reduced to ferrous iron (Fe^2+^) and exported to the cytosol (7), where ferritin acts as the major high-capacity iron storage protein (8). Interestingly, accumulations of ferritin and other “ferruginous” complexes enclosed in vacuoles were observed in intestinal epithelial cells of *D. rotundus* (9, panel D). Loss of *REP15* likely enhances iron accumulation in intestinal epithelial cells and shedding of these cells boosts iron excretion in *D. rotundus*. (D) Light microscopy image, reproduced from Figure 3 in ^13^, showing a longitudinal section of the upper villus half from the *D. rotundus* ileum. Prussian blue staining that indicates iron demonstrates the presence of iron-containing cytoplasmic granules in epithelial cells (arrow). In addition to delivering iron via the bloodstream, a macrophage-linked mechanism contributes to iron deposition in these epithelial cells ^13^. (E) Prussian blue-positive granules are present in the forming feces of *D. rotundus*, showing that these bats excrete iron by shedding iron-containing intestinal cells. The figure is reproduced from Figure 8 in ^13^.

*REP15* is specifically expressed in the gastrointestinal tract ^48^ (Figure 3B). Cellular overexpression of *REP15* decreases the amount of iron-transporting transferrin receptors on the cell surface ^46^ (Figure 3C). Since transferrin receptor availability is a limiting factor for cellular iron uptake ^49^, *REP15* likely inhibits the uptake of iron from the bloodstream into gastrointestinal cells. Consistent with this inhibitory effect, the downregulation of *REP15* in colorectal cancer cells coincides with increased intracellular iron levels in these cells ^50,51^.

Why is a gene that inhibits cellular iron uptake specifically lost in *D. rotundus?* We propose that *REP15* loss represents a strategy to boost iron excretion and thus helps vampire bats to reduce systemic iron levels. Loss of *REP15* is expected to enhance the accumulation of iron in cells of the gastrointestinal tract, where it is specifically expressed. Because the intestinal epithelium has a relatively fast turnover time ^52^, iron is subsequently eliminated by shedding iron-containing gastrointestinal cells. This general mechanism has been recently suggested as an important factor controlling iron loss in mammals ^53^.

Intriguingly, a 1980 study on the distribution of iron in the gastrointestinal tract of *D. rotundus* made observations that precisely match this hypothesis ^13^. Using histochemical staining techniques along with electron microscopy and radiography, large accumulations of iron in ferritin-containing vacuoles were identified in gastrointestinal epithelial cells that frequently desquamated into the intestinal lumen ^13^ (Figure 3D,E). This observation is consistent with enhanced cellular iron uptake in gastrointestinal cells of vampire bats to which *REP15* loss could contribute. In addition to enhanced excretion, a previous study found that vampire bats limit gastrointestinal iron absorption by increased expression of hepcidin, a factor that inhibits intestinal iron absorption ^47^. Furthermore, iron-storing ferritin genes are expanded in the common vampire bat genome ^21^ and these bats have high levels of iron-binding RFESD in the serum proteome ^54^. Thus, limited iron absorption (mediated by increased hepcidin expression), a higher capacity for iron storage (mediated by ferritin and RFESD), and enhanced iron excretion (mediated by inactivating the inhibitory factor *REP15*) help vampire bats to cope with their iron-rich diet.

### Losses of *FFAR1* and *SLC30A8* and reduced insulin secretion

*FFAR1* (free fatty acid receptor 1) encodes a G protein-coupled receptor that is highly expressed in pancreatic beta cells and senses medium to long chain free fatty acids ^48,55^. Free fatty acids augment glucose-stimulated insulin secretion and *FFAR1* is a major factor that mediates this effect ^55,56^ (Figure 4). Glucose-stimulated insulin secretion involves an initial, rapid phase, which appears to be largely dependent on the depletion of an existing pool of membrane-docked insulin secretory granules, and a second, prolonged phase, which requires the production of new granules to supply the ongoing secretion process ^55,57^. Deletion of *FFAR1* in mice reduces the amplifying effect of free fatty acids in the prolonged phase by ~50% ^56,58^. Consistent with this, *FFAR1* expression levels in human islets positively correlate with insulin secretion, which suggested that *FFAR1* deficiency could lower the insulin secretory capacities of beta cells and thus contribute to the development of type 2 diabetes ^59,60^. In addition to directly stimulating insulin secretion in beta cells, FFAR1 also amplifies insulin secretion indirectly ^55^. First, *FFAR1* is expressed in enteroendocrine cells, where its activation triggers the release of incretin hormones that stimulate pancreatic insulin secretion ^61^. Second, *FFAR1* is expressed in the brain and likely functions as a lipid sensor that influences insulin secretion through innervation of pancreatic islets ^62^. Hence, *FFAR1* amplifies insulin secretion via multiple mechanisms.

**Figure 4:**
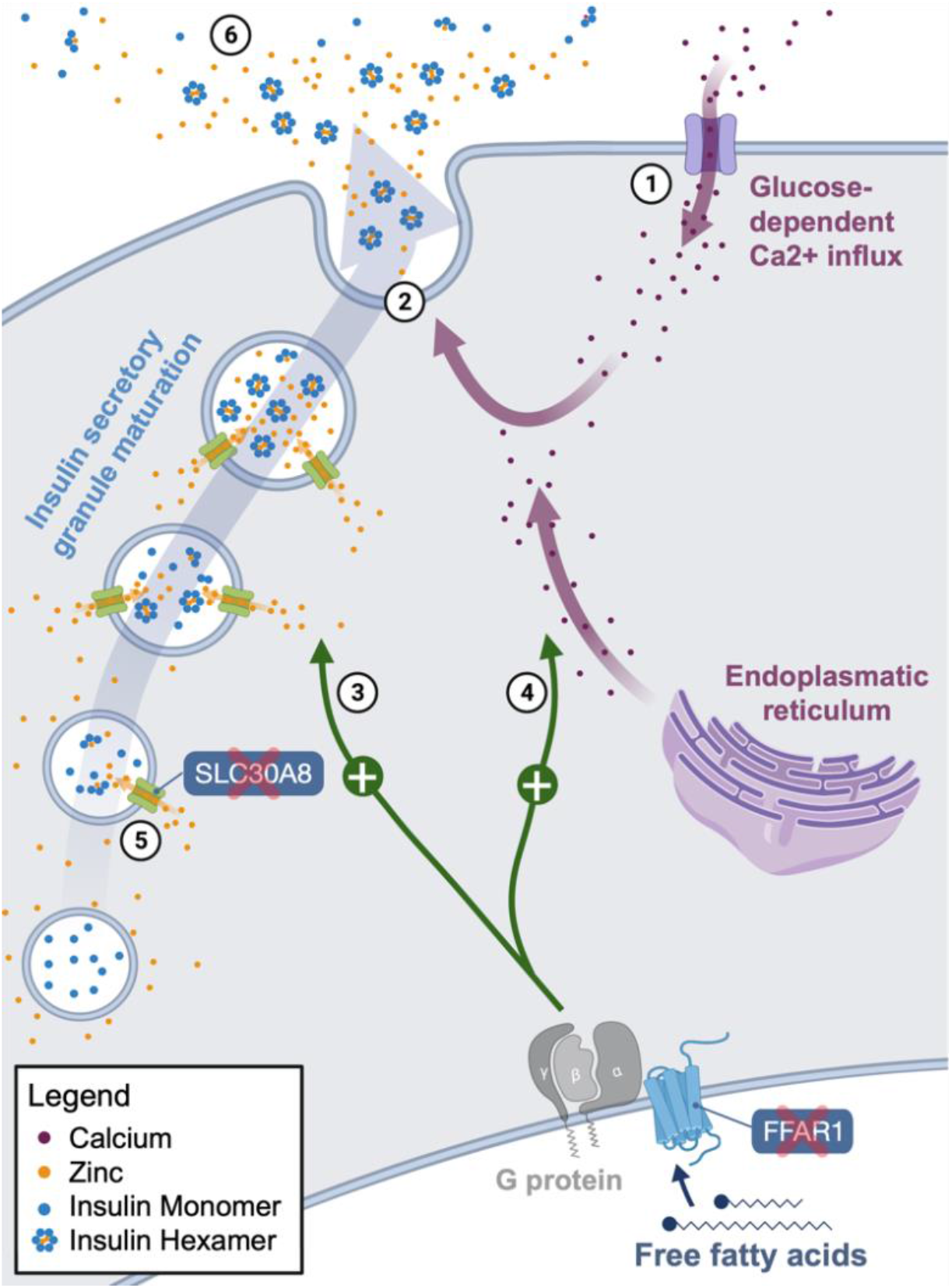
Losses of *SLC30A8* and *FFAR1* in *Desmodus rotundus* relate to reduced insulin secretion. Illustration of the roles of both genes in insulin synthesis and secretion in pancreatic beta cells. Glucose-stimulated insulin secretion is initiated by the opening of voltage-gated calcium channels (1) ^55^. The resulting influx of calcium enhances insulin secretion by stimulating exocytosis of insulin secretory granules (2). *FFAR1* encodes a G protein-coupled receptor that upon activation enhances the prolonged insulin secretion phase via (F)-actin remodeling (3). FFAR1 also triggers the release of calcium from the endoplasmic reticulum (4), which adds to the glucose stimulated calcium influx and amplifies exocytosis of secretory granules. SLC30A8 transports zinc into maturing insulin secretory granules, where zinc is essential for the formation of insulin hexamers (5). Upon secretion, these hexamers dissociate, releasing bioactive insulin monomers and zinc into the circulation (6). Loss of *FFAR1* and *SLC30A8* in *D. rotundus* is likely related to vampire bats’ reduced insulin secretion.

The free fatty acid sensor FFAR1 likely became dispensable for vampire bats for two reasons. First, the FFAR1 ligand (fatty acids) is barely present in their blood diet. Second, the low dietary sugar content likely abolishes the need for a normal glucose-stimulated insulin secretion, including the prolonged insulin secretion phase to which FFAR1 substantially contributes. Indeed, vampire bats exhibit lower basal insulin levels than other mammals and show a substantially reduced insulin secretion with prolonged hyperglycemia upon an experimental glucose overload ^10,11,16^. While having similarities to the defective insulin secretion response in human type 2 diabetes patients, in vampire bats, this is likely an adaptation to prevent hypoglycemia, since glucose must remain in the bloodstream. Whereas a previous study detected signatures of positive selection in *FFAR1* and interpreted it as a means to improve glucose utilization in *D. rotundus* ^21^, our analysis reveals that *FFAR1* is actually lost and suggests that this gene loss relates to reduced insulin secretion under normal conditions and the inability to cope with high glucose intake when experimentally challenged.

*SLC30A8* (solute carrier family 30 member 8) encodes the most highly expressed zinc transporter in pancreatic beta cells ^63,64^. In secretory granules of beta cells, zinc is required for the formation of zinc-insulin hexamers (Figure 4), which protect insulin from degradation ^65,66^. Therefore, loss of the zinc transporter *SLC30A8* may have been a consequence of or may contribute to the decreased insulin secretion phenotype of *D. rotundus*.

Another, not mutually exclusive hypothesis is that *SLC30A8* loss in vampire bats may be beneficial by reducing the requirements for body zinc levels. Even though sufficient dietary zinc is available from mammalian blood ^67,68^, it might not be efficiently absorbed by vampire bats, since high dietary iron concentrations can inhibit zinc absorption ^69^ and zinc in turn stimulates iron absorption in intestinal cells ^70^. Since blood has a high iron:zinc ratio of 66:1 ^67^, high dietary iron levels could induce zinc deficiency in vampire bats. Interestingly, *SLC30A8* knockout in mice causes extremely low zinc concentrations in beta cells ^71^. Similarly, “natural *SLC30A8* knockouts” in mammalian herbivores such as guinea pig, sheep and cow that also lost this gene ^72^ are associated with low pancreatic islet zinc concentrations ^73–75^. Since beta cells are normally among the cell types with the highest zinc concentration ^76^, it is possible that these herbivores may have lost *SLC30A8* because vegetarian diets are generally more limited in zinc content ^77,78^. Repeated loss of *SLC30A8* in the vampire bat and in herbivorous mammals could therefore represent a strategy to restrict zinc utilization to other essential physiological functions. A precondition for this strategy would be that insulin stability and secretion does not depend on zinc anymore, as shown in guinea pigs, or that insulin secretion would not be essential anymore, as it seems to be the case for vampire bats.

### Loss of *PPP1R3E* and impaired glycogen metabolism

*PPP1R3E* (protein phosphatase 1 regulatory subunit 3E) encodes a regulatory subunit of protein phosphatase 1 (PP1) ^79^. PP1 plays a central role in regulating a switch between synthesis and breakdown of glycogen, the major short-term storage form of glucose ^80^. By dephosphorylating glycogen synthase, PP1 activates this enzyme and promotes glycogen synthesis ^81^. By dephosphorylating glycogen phosphorylase, PP1 inhibits this enzyme and consequently glycogen breakdown ^81^ (Supplementary Figure 3A). The activity of PP1 depends on a regulatory subunit, which is encoded by seven different genes with different expression patterns (*PPP1R3A* - *PPP1R3G*) ^79^. These regulatory subunits are critical for PP1 activity as overexpression of *PPP1R3A*, *PPP1R3B*, *PPP1R3C*, or *PPP1R3G* in cellular or animal models increases glycogen content, whereas their knockout reduces glycogen content ^82–86^. Even though no animal studies of *PPP1R3E* exist, the gene is transcriptionally regulated by insulin and PPP1R3E binds to glycogen ^79^. Thus, *PPP1R3E* most likely functions like the other glycogen-targeting subunits. With the exception of *PPP1R3E* loss in *D. rotundus*, none of the seven regulatory subunit-encoding genes exhibit inactivating mutations in any analyzed bat (Supplementary Figure 3B).

The specific loss of *PPP1R3E* likely relates to the low glycogen concentrations in *D. rotundus* ^17^. In fact, fed vampire bats have hepatic glycogen stores that are ~85% smaller than in fruit-eating bats and ~60% smaller than in other mammals fed on high-protein diets ^17^. Since sufficient glycogen stores are important to withstand periods of fasting, loss of *PPP1R3E* and the associated smaller glycogen stores may also contribute to the observed starvation vulnerability of vampire bats.

### Loss of *CTSE* and altered stomach function

The loss of *CTSE* (cathepsin E) in *D. rotundus* may be a consequence of extensive morphological and physiological modifications of their stomach, which is unparalleled by any other mammalian species ^12,87^. Most notably, the stomach of *D. rotundus* experienced fundamental remodeling from a compact muscular organ involved in mechanical and chemical digestion towards a distensible structure that functions primarily to store large amounts of ingested blood and serves as a major site of fluid absorption ^12,87,88^ (Supplementary Figure 4). CTSE is an intracellular protease that is highly expressed in gastrointestinal tissues, particularly in the stomach ^48,89^. In the stomach, CTSE protein normally localizes to the canaliculi (invaginations) of parietal cells, which secrete hydrochloric acid, indicating that CTSE might be involved in secretory processes ^90^. In *D. rotundus*, the canaliculi of parietal cells are specifically enriched in iron ^13^, suggesting that their gastric parietal cells experienced a functional shift from hydrochloric acid secretion towards participating in iron excretion, which could have rendered CTSE dispensable. Consistent with this, *CTSE* has been reported lost in platypus ^91^, which also lacks an acid-secreting stomach. However, *CTSE* is also lost in Cetartiodactyla that possess acid-secreting stomachs ^92^ and, in addition to a possible role in gastric acid secretion, *CTSE* also has immune-related functions ^89^. Thus, other *CTSE* functions could have also contributed to or represent the primary reason of *CTSE* loss in *D. rotundus*.

### Loss of *ERN2* and low dietary fat content

The loss of *ERN2* (endoplasmic reticulum to nucleus signaling 2) in *D. rotundus* is likely a consequence of the low fat content of their blood diet. *ERN2* is specifically expressed in gastrointestinal epithelia ^93^ and encodes a transmembrane protein that inhibits the production of chylomicrons, which are lipoprotein particles that transport dietary lipids absorbed in enterocytes to other tissues ^94,95^. Enterocytes of *ERN2* knockout mice were shown to secrete more chylomicrons, resulting in increased lipid absorption and a more pronounced hyperlipidemia on high-fat diets ^94,95^. Interestingly, on a standard diet, *ERN2* knockout mice did not exhibit hyperlipidemia, indicating that *ERN2* limits intestinal lipid absorption only under conditions where excessive fat is available. Since blood has a very low fat content ^11^, it is conceivable that regulatory mechanisms, which normally limit lipid absorption, became dispensable during vampire bat evolution, resulting in loss of *ERN2*.

### Loss of the pancreatic chymotrypsin *CTRL*

Pancreatic proteases such as chymotrypsinogens and trypsinogens are among the most essential digestive enzymes ^96^. They are produced by acinar cells of the exocrine pancreas and are secreted as inactive zymogens which, upon activation in the small intestine, digest proteins ^96^. *CTRL* (chymotrypsin like) encodes one of four chymotrypsinogen isozymes and is considered a predictive biomarker for human pancreatic cancer ^97^. In mice, CTRL represents a minor chymotrypsin isoform, constituting ~10% of the chymotrypsinogen pool ^98^. *In vitro* experiments showed that CTRL cleaves trypsinogens and thus inhibits the activation of protein-digesting trypsin ^98^. Consistently, *CTRL* knockout in mice leads to a reduced activation of chymotrypsins and a slightly higher trypsin activity ^98^, which could be relevant for the unique, protein-rich diet of vampire bats. Interestingly, recent evidence also indicates that acinar proteases including CTRL contribute to proliferation of beta cells ^99^, providing another possible explanation for the loss of this gene in a bat species that exhibits a reduced beta cell mass ^16^.

### Loss of *CYP39A1* and advanced social behavior

The loss of *CYP39A1* (cytochrome P450 family 39 subfamily A member 1) could have contributed to the evolution of vampire bat’s exceptional social behaviors and cognitive abilities ^20,100,101^. *CYP39A1* encodes an oxysterol 7-α-hydroxylase enzyme ^102^. Apart from a minor contribution to hepatic bile acid synthesis (Supplementary Figure 5), CYP39A1 is the only enzyme expressed in the brain that can degrade the cholesterol metabolite 24S-hydroxycholesterol ^103–106^. 24S-hydroxycholesterol is an intermediate by-product during cholesterol elimination from the brain ^107^. Unlike cholesterol, 24S-hydroxycholesterol can pass the blood-brain barrier. Therefore, the loss of the 24S-hydroxycholesterol-degrading CYP39A1 is expected to result in elevated systemic levels of this metabolite, which was observed in humans with *CYP39A1* loss-of-function alleles ^108^.

Since 24S-hydroxycholesterol also plays important neurophysiological roles, we hypothesize that *CYP39A1* loss may be connected to the exceptional cognitive and social capabilities of *D. rotundus*. 24S-hydroxycholesterol is a potent allosteric activator of *N*-methyl-D-aspartate receptors (NMDAR), which are glutamate-gated ion channels that mediate synaptic plasticity and memory formation ^109,110^. Consistent with a positive effect of 24S-hydroxycholesterol on NMDAR activity and cognitive function, increased cerebral 24S-hydroxycholesterol levels improve spatial memory function in mice ^111,112^, whereas reduced levels cause impaired learning and memory function ^113,114^. NMDAR activity also exerts a strong influence on social behavior, as NMDAR agonists improve social behavior in rodents, whereas antagonists impair it ^115^.

Intriguingly, vampire bats, the only lineage in our dataset that lost the 24S-hydroxycholesterol-metabolizing *CYP39A1* gene, are distinguished from other bats by their exceptional social behavior and cognitive abilities. They reciprocally share regurgitated blood with roost mates that failed to obtain a nightly meal and would otherwise face starvation ^19^. The decision with whom to share blood is primarily driven by recognizing contact calls of individuals that provided past blood donations, which demonstrates exceptional long term social memory ^20,100^. Furthermore, non-kin adoption of orphaned offspring occurs in vampire bat colonies ^101^. Vampire bats also form long-lasting social bonds, evident by observations that individuals who cooperated in captivity retain their social network when released into the wild ^116^. Vampire bats, as perhaps the most socially-advanced bats, were also found to have the largest relative neocortical volume among 276 measured bat species ^117^. Interestingly, the 24S-hydroxycholesterol-interacting subunit of NMDAR (*GLUN2B*) is highly expressed at birth and plays an important role in cortical development ^110,118^, raising the possibility that *CYP39A1* loss may also be connected to the large neocortical volume of *D. rotundus*.

### Loss of cone phototransduction genes suggests rod monochromacy

Our strict genomic screen revealed that *PDE6H* is exclusively lost in vampire bats. *PDE6H* encodes the γ’-subunit of the cone phosphodiesterase, a component of the phototransduction cascade in cone cells ^119^. Loss of function mutations in *PDE6H* cause total color blindness in humans ^119^, but not in mice, where the rod phosphodiesterase isoform *PDE6G* compensates for *PDE6H* loss ^120^. Given this cross-species variability, we investigated additional genes required for phototransduction in rods and cones in the 26 focal bat species. For rods, we found no phototransduction component (*RHO*, *GNAT1*, *GNB1*, *GNGT1*, *PDE6A*, *PDE6B*, *PDE6G*) to be lost. Among the genes required for cone phototransduction (*OPN1SW*, *OPN1LW*, *GNAT2*, *GNB3*, *GNGT2*, *PDE6C*), we confirmed previously-reported losses of the short-wavelength sensitive opsin (*OPN1SW*) in the vampire bat and six other bats ^121–124^ (Supplementary Figure 6). In addition, our analysis detected that *PDE6C* is lost in the vampire and four other bats (Supplementary Figures 6, 7). Inactivating mutations in *PDE6C* in both humans and mice abolish cone function entirely, resulting in rod monochromacy, a condition where only rod photoreceptors remain functional ^125,126^.

While ancestral Chiroptera were inferred to have two types of cones and dichromatic vision, many bats, including *D. rotundus*, lack cones that express a functional short-wavelength sensitive opsin (*OPN1SW*) ^121–124^. Whereas the lack of a functional *OPN1SW* alone suggested color blindness but maintenance of functional cones expressing the conserved *OPN1LW* (long wave sensitive opsin) gene, our discovery of repeated *PDE6C* losses suggests that *D. rotundus* and four other nocturnal Noctilionoidea may be functional rod monochromats. This condition, where all cone-based photoreception is abolished even if vestigial cone cells can still be detected, has been suggested for only a few other mammalian lineages so far ^127^. In light of their ecology, rod monochromacy is plausible in vampire bats, which lost both *PDE6C* and *PDE6H*. These strictly nocturnal bats are most active during the darkest periods of the night and even avoid moonshine ^128^. Furthermore, to locate their prey, vampire bats have evolved non-visual sensory adaptations such as highly specialized auditory adaptations ^129^, and an infrared sensing capability, which is unique among mammals ^6^.

### Loss of the immune-related *RNASE7* and a different pathogen profile of blood

Feeding exclusively on blood, the immune system of vampire bats is regularly challenged with blood-borne pathogens ^130^. However, because blood exhibits a low bacterial abundance but different species composition ^131,132^, their intestinal tract is likely exposed to a different diversity of pathogens. We discovered that the immune-related *RNASE7* (ribonuclease A family member 7) gene is lost in *D. rotundus* and intact in all other analyzed bats.

*RNASE7* encodes a secreted ribonuclease that has potent antimicrobial activity against various microorganisms ^133^. In humans, *RNASE7* is highly expressed in most epithelia including skin, urothelium and respiratory tract epithelium ^48,133^. The gene is significantly downregulated in the skin and urinary tract of diabetic patients (potentially because *RNASE7* expression can be induced by insulin signaling), which reduces innate immune defense capacities and likely contributes to the much higher incidence of bacterial skin and urinary tract infections in diabetic patients ^134,135^. The exclusive loss of *RNASE7* in vampire bats, which might also be related to their reduced insulin secretory capacities, predicts that vampire bats have a reduced capacity for secreting bactericidal peptides. While a previous study found that *RNASE7* evolved under positive selection in vampire bats, which was interpreted as an adaptation towards increased exposition to blood-borne pathogens ^21^, our analysis shows that *RNASE7* is clearly inactivated, raises the possibility that its loss may be a consequence of exposure to a different pathogen diversity.

## Discussion

Here, we used long read (HiFi) sequencing to generate a haplotype-resolved genome assembly of high completeness, contiguity and base accuracy for the common vampire bat *Desmodus rotundus*. Adding to previous studies ^34,136^, our side-by-side comparison of gene content and inactivating mutations demonstrates that the HiFi assembly not only improves gene completeness but also base accuracy. A haplotype-resolved assembly also facilitates discriminating between homozygous and heterozygous (inactivating) mutations, which is more difficult in collapsed diploid assemblies. Using this new *D. rotundus* assembly and existing genomes of 25 other bats, we performed a genome-wide screen for genes that are specifically lost in the vampire bat lineage. This screen revealed 10 previously-unknown gene losses that are likely associated with derived phenotypic features of vampire bats, such as reduced insulin secretion (*FFAR1*, *SLC30A8*) and glycogen synthesis (*PPP1R3E*), a distinct gastric physiology (*CTSE*), and exceptional social behavior and cognitive abilities (*CYP39A1*). The loss of other genes (*ERN2*, *CTRL*, *REP15*) is likely related to the biased nutrient composition of blood, which features low fat, high protein and high iron contents. Consistent with an association to vampire bat-specific changes in metabolism and digestion, many of these genes are highly expressed in relevant organ systems such as the pancreas (*FFAR1*, *SLC30A8*, *CTRL*) and gastrointestinal tissues (*REP15*, *CTSE*, *ERN2*). Finally, gene loss can also indicate previously-unknown phenotypes, exemplified here by the loss of *PDE6H* and *PDE6C*, which suggest the complete lack of cone photoreceptor function (functional rod monochromacy). Similarly, the loss of *RNASE7* may indicate differences in the immune system between *D. rotundus* and other bats, which deserves further studies. In particular, while bats are generally known for tolerating a larger diversity of viral and other pathogens, it is conceivable that their specialized diet exposes *D. rotundus* to a distinct pathogen profile, largely constrained to blood-borne pathogens. This hypothesis could be tested using metagenomics or related technologies to systematically characterize and compare pathogen profiles of vampire and other bats, ideally in matched environments.

Our finding that vampire bats, which naturally exhibit a reduced capacity for insulin secretion ^16^, have lost *FFAR1*, a beta cell receptor that stimulates insulin secretion, provides an interesting contrast to fruit eating bats of the phyllostomid and pteropodid families, which exhibit an increased capacity for insulin secretion.

These fruit bats have convergently adapted to sugar-rich diets ^137,138^ and have convergently lost *FFAR3* ^23,26^, a different beta cell receptor that inhibits insulin secretion ^139^. This suggests that losses of two different genes with opposite effects on insulin secretion are involved in opposite phenotypes in bats.

Which genes are lost during the course of evolution is influenced by several factors. One important factor underlying gene loss, including those identified here, is the absence of selection to preserve the function of certain genes as a consequence of adaptation to a new environment or different lifestyle. However, despite dispensability of gene function, several other factors constrain which genes are permitted to be lost in evolution ^25^. For example, genes lost in mammals are generally depleted in essential functions and tend to have a lower degree of pleiotropy ^92^. Indeed, the few known examples of pleiotropic gene losses in mammals refer to cases where most or all functions of the gene became dispensable (e.g. *KLK8*, *INSL5*, *RXFP4*, *SLC4A9* ^28,29,140^) or where additional gene functions are compensated by functionally related proteins (e.g. *ACOX2, SLC27A5* ^141^). Consistent with pleiotropy being a key restricting factor, the gene loss cases detected here largely refer to genes with specific functions and a low or no degree of pleiotropy. In addition to pleiotropy and essentiality, restricted expression patterns appear to be another factor permitting gene loss. For example, while the inactivated gene *ERN2* exhibits a specific expression in the gastrointestinal tract, the ubiquitously expressed and functionally similar *ERN1* gene ^93^ is intact in the vampire bat genome. Similarly, *REP15*, *SLC30A8*, *CTSE*, and *CTRL* have tissue-restricted expression patterns.

While dispensability of gene function is certainly the main explanation for gene losses in the vampire bat lineage, loss of ancestral genes can sometimes be beneficial in evolution and contribute to adaptations ^25,26,28,29,142^. For example, the loss of *REP15* could have contributed to enhanced iron elimination through shedding of iron-containing gastrointestinal cells, which likely represents an adaptation of vampire bats to their iron-rich blood diet. Another example is *CYP39A1*, whose loss is expected to result in elevated systemic levels of 24S-hydroxycholesterol. Since elevated levels of this cholesterol metabolite can positively affect cognitive function in animal models ^111,112^, it is conceivable that loss of *CYP39A1* in the vampire bat lineage may have contributed to their exceptional social behavior and cognitive abilities, which is a unique feature among bats.

In summary, our study provides novel insights into the genomic changes related to adaptations to sanguivory and the evolutionary importance of gene losses in general. Notably, a comprehensive understanding of how sanguivory, as a unique dietary specialization, has evolved, requires not only high-quality genomic resources but also data on the organismal biology of vampire bats. Our study reveals gaps in our knowledge of vampire bat physiology, metabolism and immunity, highlighting the need to better characterize the phenotypic side of adaptations to their unique diet.

## Methods

### Sample collection for sequencing

A *D. rotundus* male individual (ROMM126221) was collected in a mist net at Sabajo, Para, Suriname on 26 September 2017 as part of a larger environmental baseline study and euthanized with an overdose of isoflurane. The research permit was approved on 8 June 2017 by the Nature Conservation Division, Suriname Forest Service. Animal Use Protocol #2017-19 was approved by the Animal Care Committee of the Royal Ontario Museum. The use of wild mammals in research followed the guidelines of the American Society of Mammalogists ^143^.

### DNA extraction, library preparation and whole genome sequencing

Sampled tissues were snap-frozen in liquid nitrogen immediately after dissection and stored at −80 °C until further processed. Mixed tissue of heart, liver, and spleen (23 mg) was used for high molecular weight DNA extraction using the Nanobind Tissue Big DNA Kit (Circulomics, MD) according to the manufacturer’s instructions. Final DNA purity and concentrations were measured using NanoPhotometer^®^ (Implen GmbH, Munich, Germany) and Qubit Fluorometer (Thermo Fisher Scientific, Waltham, MA). Two SMRTbell libraries were constructed following the instructions of the SMRTbell Express Prep kit v2.0 (Pacific Biosciences, Menlo Park, CA). The total input DNA for each library was approximately 5 μg. The libraries were loaded at an on-plate concentration of 80 pM using diffusion loading. Four SMRT cell sequencing runs were performed on the Sequel System IIe in CCS mode using 30 hour movie time with 2 hours pre-extension and sequencing chemistry V2.0. The Dovetail Omni-C library was prepared from mixed tissue (heart, liver, and spleen) using the Dovetail^™^ Omni-C^™^ kit (Dovetail Genomics, Scotts Valley, CA, USA) following the manufacturer’s protocol (manual version 1.2 for mammalian samples). The Omni-C library was sequenced on a NovaSeq 6000 platform at Novogene (UK), generating 400 million 2 × 150 bp paired-end reads totaling 120 Gbp. The fragment size distribution and concentration of the final PacBio and Dovetail Omni-C library were assessed using a TapeStation (Agilent Technologies) and a Qubit Fluorometer (Thermo Fisher Scientific, Waltham, MA), respectively.

### *D. rotundus* genome assembly

We called CCS reads (rq > 0.99) from the subreads.bam file using PacBio ccs (version 6.0.0, https://github.com/PacificBiosciences/ccs). We then created the two contig assemblies using hifiasm (version 0.15.1-r331) ^144^ with arguments −l2 --h1 --h2 using the ccs reads and the HiC reads as input. This resulted in one contig assembly for each of the haplotypes in this species, which both went independently into the following scaffolding, gap-closing and polishing steps.

For scaffolding, we used SALSA2 (git commit 1b76bf63efb973583647a1eb95863d33ee6e09ad) ^145^ and the Dovetail Omni-C mapping pipeline (https://omni-c.readthedocs.io/en/latest/fastq_to_bam.html). Briefly, we mapped HiC reads using BWA-MEM (version 0.7.17-r1188) ^146^. Alignments were filtered using pairtools (version 0.3.0) with arguments parse --min-mapq 40 --walks-policy 5unique --max-inter-align-gap 30 to retain those alignments with high mapping quality, the most 5’ alignments for conflicts and alignments without large gaps. We removed potential PCR duplicates using pairtools dedup. The resulting read-sorted bed file was used as input for SALSA2. We then performed a number of manual curation rounds to correct scaffolding errors and to scaffold those contigs which were not automatically scaffolded. To this end, we used cooler (version 0.8.11, https://github.com/open2c/cooler) and HiGlass ^147^ to visually inspect the HiC maps and re-scaffolded the assembly using SeqKit (version 0.13.2) to re-arrange contigs and scaffolds into chromosome-level scaffolds.

After scaffolding, we closed additional assembly gaps using the lower-quality PacBio CLR read data that did not yield a CCS read. To this end, we mapped the original subreads.bam files to the scaffolded assembly using pbmm2 (version 1.3.0) with arguments --preset SUBREAD −N 1. Based on the read-piles created by reads spanning across gap regions, we computed consensus sequences for the gap regions and their 2 kb up/downstream flanks using gcpp (version 2.0.2-2.0.2). We filtered for high-confidence consensus sequences by requiring that no N’s and no lower-case (acgt) characters remain in the consensus sequence. We then replaced the assembly gap and flanking region with high-confidence consensus sequences.

Despite the high accuracy of PacBio CCS reads, an assembly can still contain base errors due to errors in the consensus sequence calculation. In addition, lower base accuracy likely exists in the closed gap regions. To improve base accuracy, we polished both haplotype assemblies using all CCS reads. We tested both freebayes (version 1.3.2, https://github.com/freebayes/freebayes) and DeepVariant (version 1.1.0) ^148^. Based on numbers of genes with inactivating mutations (computed by TOGA, see below) and the QV values computed by Merqury (version 1.0) ^32^, DeepVariant vastly outperformed freebayes in correcting base errors. Therefore, for the final polishing round, we mapped all CCS reads to the scaffolded, gap-closed assemblies using pbmm2 with arguments: --preset CCS −N 1 and called variants using DeepVariant. We then filtered for sites with ‘genotype 1/1’ to specify that all or nearly all reads support an alternative sequence at this position and a ‘PASS’ filter value to specify that the site passed DeepVariant’s internal filters. We then corrected base errors using bcftools consensus (version 1.12). Importantly, this procedure does not ‘correct’ any heterozygous or polymorphic regions but only those that are incorrect and not supported by any CCS reads.

### Repeat masking

We used RepeatModeler (http://www.repeatmasker.org/, parameter -engine ncbi) to generate a *de novo* repeat library for the new *D. rotundus* genome assembly and used the resulting library with RepeatMasker (version 4.0.9, parameters -engine crossmatch -s) to soft-mask the genome.

### Genome alignments

We used the human hg38 genome assembly as the reference and considered *D. rotundus* and 25 other bats as query species (Supplementary Table 1). To generate pairwise genome alignment chains as input for TOGA ^33^, we first used LASTZ (version 1.04.03) ^149^ with parameters (K = 2400, L = 3000, Y = 9400, H = 2000 and the lastz default scoring matrix). These parameters are sensitive enough to capture orthologous exons between placental mammals ^150^. Then, we used axtChain ^151^ (default parameters except linearGap=loose) to compute co-linear alignment chains, RepeatFiller ^152^ (default parameters) to capture previously-missed alignments between repetitive regions and chainCleaner ^153^ (default parameters except minBrokenChainScore=75000 and -doPairs) to improve alignment specificity.

### Using TOGA to infer orthologous genes and detect gene losses

To compare gene completeness and to screen for gene losses, we used TOGA ^33^, a method that uses pairwise genome alignment chains between an annotated reference genome (here human hg38 assembly) and other query species. Briefly, TOGA uses machine learning to infer orthologous loci for each reference transcript, utilizing that orthologous genes display more alignments between intronic and flanking intergenic regions ^33^. TOGA then projects each reference transcript to its orthologous query locus using CESAR 2.0 ^154^, a Hidden Markov model method that takes reading frame and splice site annotation of the reference exons into account. CESAR avoids spurious frameshifts, is able to detect evolutionary splice site shifts and precise intron deletions ^154,155^. Using the CESAR alignment, TOGA determines whether the transcript has inactivating mutations (frameshifting mutations, premature stop codons, splice site disrupting mutations, deletions of entire coding exons). TOGA also uses orthologous alignment chains to detect genes that are entirely deleted in the query assembly and distinguishes real exon or gene deletions from missing sequence caused by assembly gaps.

TOGA classifies each transcript into one of five categories. Classification ‘intact’, ‘partially intact’ and ‘missing’ refers to transcripts that lack inactivating mutations in the middle 80% of the coding region. TOGA specifically considers this central part of the coding region, since truly conserved genes can have inactivating mutations in the first or last 10% of the coding region (near the N- or C-terminus) ^33,155^. For intact transcripts, the central part of the coding region is completely present in the assembly, whereas for partially intact (≥50% present) and missing (<50% present) transcripts some coding parts are missing due to assembly gaps or fragmentation. Classification ‘lost’ refers to transcripts that have at least two inactivating mutations in at least two exons. All other transcripts, having a single inactivating mutation or mutations in only a single exon, are classified as ‘uncertain loss’, reflecting the possibility that only an exon but not the entire gene could be lost.

Considering all input transcripts of a gene, TOGA uses the precedence order ‘intact’, ‘partially intact’, ‘uncertain loss’, ‘lost’, ‘missing’ to provide a gene classification. That means, TOGA only classifies a gene as lost if all its transcripts are classified as lost (at all orthologous loci if there is more than one).

### Gene completeness among bat genomes

To compare gene completeness between assemblies, we first obtained a set of genes that likely already existed in the placental mammal ancestor. To this end, we used the human GENCODE V38 (Ensembl 104) gene annotation ^156^ as input and applied TOGA to the genomes of 11 afrotherian (represented by aardvark, cape golden mole, small Madagascar hedgehog, Talazac’s shrew tenrec, cape elephant shrew, dugong, manatee, Asiatic elephant, African savanna elephant, cape rock hyrax, yellow-spotted hyrax) and five xenarthran (represented by southern two-toed sloth, Hoffmann’s two-fingered sloth, nine-banded armadillo, giant anteater, southern tamandua) mammals. Since each input gene is by definition present in human (superorder Boreoeutheria), we considered a gene as ancestral if it is also conserved in the other two basal placental superorders Afrotheria and Xenarthra. Specifically, we selected genes that are classified by TOGA as intact or partially intact in at least one afrotherian and at least one xenarthran genome. Note that this requirement assures placental mammal ancestry irrespective of the exact basal split of placental mammals, which is difficult to resolve ^157^. This procedure resulted in a set of 18,430 genes.

Next, we applied TOGA to *D. rotundus* and the 25 other bat genomes (Supplementary Table 1). Considering the 18,430 ancestral genes, we counted per species how many gene have an intact reading frame (TOGA classification ‘intact’), have inactivating mutations (TOGA classifications ‘loss’ and ‘uncertain loss’) or have missing sequence (TOGA classifications ‘partially intact’ and ‘missing’). This breakdown is shown in Figure 1C.

### Screen for gene losses

To identify gene losses that among bats occurred specifically in the vampire bat lineage, we used the above-generated TOGA data for the 26 bats. We first filtered for genes that have at least one inactivating mutation in the middle 80% of the coding sequence (TOGA classification ‘lost’ or ‘uncertain loss’) in both *D. rotundus* haplotype assemblies.

Base errors can also occur in other bat assemblies, which may result in missing real vampire bat-specific gene losses (false negatives). To reduce the false negative rate, we conducted the initial screen with less strict requirements and allowed per gene up to three non-vampire bat species to be classified as ‘missing’, ‘partially missing’ or ‘uncertain loss’, resulting in 52 candidate genes. Then, to assess which of these candidate genes are a true vampire specific gene losses, we mapped available Illumina reads of the same species to each included genome assembly using BWA-MEM (version 0.7.7-r441) ^146^ and removed duplicates with Picard (version 2.21.4) (http://broadinstitute.github.io/picard) (SRA identifiers are listed in Supplementary Table 4) and examined whether putative inactivating mutations are supported by raw reads and if they are heterozygous or homozygous. We excluded all genes as non-vampire specific losses that have homozygous mutations in at least one other bat species and furthermore required that not more than 60% of the coding sequence remains functional in *D. rotundus*. We kept *SLC30A8*, which has a heterozygous stop codon in exon 4 in *Micronycteris hirsuta*. We excluded genes belonging to the large and fast-evolving olfactory receptor family. This procedure revealed the 13 vampire specific gene losses that are discussed in the main text.

Finally, to confirm the correctness of all 10 previously-unknown vampire bat-specific gene losses, we manually inspected the pairwise alignment chains in the UCSC genome browser ^158^, which showed that the remnants of all 10 genes are located in a conserved gene order (synteny) context. Inspecting chiropteran pairwise alignment chains also showed that none of the 10 genes exhibit duplications in *D. rotundus* or other bats, which excludes the possibility that a functional gene copy remained in *D. rotundus*. Finally, we further verified the correctness of the inactivating mutations by requiring that the gene is also classified as ‘lost’ or ‘uncertain loss’ in the Illumina *D. rotundus* assembly with at least one shared homozygous (based on aligned Illumina sequencing reads) inactivating mutation, thus providing base support from two sequencing technologies.

### Relaxed selection

To test the 10 genes for relaxation of selection, we obtained pairwise codon alignments between human and each query bat from TOGA. Codons affected by frameshifting insertions or deletions and premature stop codons were replaced by ‘NNN’ codons to maintain a reading frame. A multiple codon alignment was produced with MACSE v2 ^159^ and cleaned with HmmCleaner ^160^ with default cost values. The resulting alignments were used to investigate whether the gene evolves under relaxed selection with RELAX ^45^, specifying the *D. rotundus* branch as the foreground and all other branches as background.

### Gene expression

We investigated RNA expression of the 10 target gene losses using available transcriptome sequencing data for *D. rotundus* from 11 tissues (heart, stomach, intestine, liver, gallbladder, pancreas, spleen, kidney, olfactory epithelium, eye and tongue). The paired-end sequencing read sets were downloaded from NCBI (accessions listed in Supplementary Table 5) and mapped to the new *D. rotundus* haplotype 1 assembly using STAR (version 2.7.3) ^161^. Target loci were then manually examined in IGV ^162^.

### Whole blood iron measurements

All animal captures were approved by the National Environmental Office (License number 77322-1, Sisbio, Brazil) and all experiments were approved by the Ethics Committee from Federal University of Viçosa (License number 10/2021, CEUA/UFV, Brazil). Nine *D. rotundus*, sixteen *Artibeus lituratus* and four *Myotis myotis* individuals were captured with mist nets in Viçosa, Minas Gerais, Brazil and euthanized through cervical dislocation followed by decapitation. All blood samples (100 - 400 μl) were dried using a stove and mineralized in a nitric-perchloric acid solution (1.5 mL total volume) until organic matter was removed. The final extract was used to determine iron concentrations through atomic absorption spectrophotometry (Shimadzu AA-6701F). Whole blood iron levels were compared between species with a two-sided t-test.

## Data availability

The haplotype-resolved *D. rotundus* assembly and the TOGA annotations are freely available at https://bds.mpi-cbg.de/hillerlab/VampireBatGenome/. The sequencing data and genome assembly is also uploaded to NCBI.

## Acknowledgments

We would like to thank the UCSC genome browser group for providing software and genome annotations, and the Computer Service Facilities of the MPI-CBG and MPI-PKS for their support. We also thank the Genome Technology Center (RGTC) at Radboudumc for the use of the Sequencing Core Facility (Nijmegen, The Netherlands). Furthermore, we want to thank Robert Cornman for support concerning the *Aeorestes cinereus* assembly, and David Lagman and Dan Larhammar for helpful feedback on the phototransduction cascade. Figures 3C, 4 and S3A created with Biorender (https://biorender.com). This work was supported by the Max Planck Society, the National Council for Scientific and Technological Development (CNPq, Brazil), funding from Environmental Services & Support (ESS) for fieldwork in Suriname, and the LOEWE-Centre for Translational Biodiversity Genomics (TBG) funded by the Hessen State Ministry of Higher Education, Research and the Arts (HMWK).

